# Comparisons of a Novel Air Sampling Filter Material, Wash Buffers and Extraction Methods in the Detection and Quantification of Influenza Virus

**DOI:** 10.1101/441154

**Authors:** Terrance A. Thedell, Corey L. Boles, David M. Cwiertny, Jiajie Qian, Grant D. Brown, Matthew W. Nonnenmann

## Abstract

Quantification of aerosolized influenza virus is used for determining inhalation exposure. Several bioaerosol samplers and analytical methods have been used; however, the detection and quantification of influenza virus among aerosol samples remains challenging. Therefore, improved viral aerosol measurement methods are needed. This study evaluated influenza virus recovery among three filter types polytetrafluoroethylene, polyvinylchloride and polystyrene. Polytetrafluoroethylene, polyvinylchloride are fabricated filter materials and commonly used in the scientific literature to sample for viral aerosols. A novel, electrospun polystyrene filter material may improve viral aerosol recovery during filter-based air sampling. The filter materials were compared across the following conditions: treated with or without air, filter wash buffer (HBSS or PBS), and viral RNA extraction method (QIAamp Viral RNA Mini Kit or Trizol). Twenty trials were completed in a chamber and samples were analyzed using RT-qPCR. Viral recovery was significantly different (p-value < .0001) by filter type. Polystyrene filter use resulted in recovery of the most viral RNA. Air sampling did not affect the recovery of viral RNA from the filter materials (p-values > 0.05). Viral RNA concentrations were significantly different across extraction methods for all comparisons (p-values < 0.05). Our results demonstrated that the novel polystyrene filter material resulted in the highest concentration of extracted RNA compared to the commonly used polytetrafluoroethylene and polyvinylchloride, which we speculate may be related to the chemical composition of the filter material (*e.g.*, polystyrene is an aromatic hydrocarbon whereas polytetrafluoroethylene and polyvinylchloride contain more polar, and thus potentially reactive, carbon-halogen bonds). Air sampling did not have an effect on viral RNA recovery. Using Hanks Balanced Salt Solution with QIAamp Viral RNA Mini Kit, and Phosphate-buffered saline with the Trizol extraction, resulted in the most viral RNA recovery.

## Introduction

Influenza is a contagious respiratory illness caused by influenza viruses that infect the nose, throat, and lungs, causing annual epidemics, and occasional pandemics. The World Health Organization (WHO) estimates that influenza epidemics cause 3-5 million cases of severe illness and up to 500,000 deaths worldwide.^1^ Influenza viruses are enveloped negative-strand ribonucleic acid (RNA) viruses^2^, and range from 80-120 nanometers (nm) in diameter.^3,4^

Influenza viruses can be transmitted by direct contact with infected individuals, exposure to virus-contaminated objects, or inhalation of infectious aerosols.^5,6^ Little is known definitively about the modes of influenza transmission.^7^ However, influenza viruses are spread by the inhalation of infectious aerosols, with the majority of these aerosols being less than 5 μm in aerodynamic diameter.^8,9^

There are several designs available to use as air samplers for the collection of bioaerosols. A number of these samplers have been used for influenza virus detection, but the most commonly used samplers are liquid impingers, solid impactors, as well as filter-based samplers.^10,11^ However, few researchers have been able to detect aerosolized influenza viruses in a field setting. The reasons are unclear, but speculation persists about the impact of environmental factors (e.g., relative humidity) and the impact of sampling methods (i.e., filter materials) on influenza virus recovery.^10,12,13^ Samplers like the SKC Biosampler have been developed to minimize the effects of sampling stress and maximize recovery. The SKC Biosampler has been found to have greater viral recovery compared to other commonly used filter-based samplers; however, the differences across samplers were not statistically significant.^13–15^

Filters are used frequently as a sampling media in personal samplers in the field of exposure assessment. Filter based samplers have also been used in the collection of aerosolized viruses due to their efficiency in collecting sub-micrometer particles^10,16^, and ability to be used for longer periods in the field compared to impinger based methods.^17,18^ Polytetrafluoroethylene (PTFE), polyvinyl chloride (PVC), polycarbonate, and gelatin-containing filters have all been used to collect virus^10,19^, with PTFE and PVC being the most efficient among filter materials tested.^16,19^ Nevertheless, the viral recovery from PTFE and PVC filter materials remains inefficient.^10^

There are many other materials that can be used to fabricate filters for collection of bioaerosols. For example, a new filter material made from polystyrene (PS) has been developed to be used in the collection of bioaerosols. PS filters have been fabricated from electrospun fibers, yielding materials with large surface areas and small pore sizes that make them very efficient in filtering applications.^20^ The hydrophobicity of the PS filter material may assist the viral capsid attachment proteins and other biomolecules to adsorb to the fiber surfaces.^21^ Furthermore, the use of PS as a centrifuge tube material is common in the field of molecular biology, therefore it is hypothesized that PS will perform well as a filter material for sampling aerosolized virus in the field of exposure science.

After viral aerosol collection, the virus is removed from the sampling media by vortexing and using a wash buffer. The function of the wash buffer [i.e., Hanks Balanced Salt Solution (HBSS) and Phosphate-Buffered Saline (PBS)], is to maintain the pH and osmotic balance, provide water and essential inorganic ions for the target organism.^22^ Wash buffers (i.e., HBBS, PBS) have been used in previous studies.^14,15,23–28^ However, there is limited information quantifying the variability in viral recovery using different types of buffers to remove viruses from filter materials.

Once the virus is removed using the wash buffer, extraction of viral RNA is needed to detect and quantify the virus collected. Influenza virus is an RNA virus, therefore, extracting and purifying viral RNA efficiently is critical for accurate detection and quantification. The QIAamp Viral RNA Mini Kit and Trizol method have been have been extensively used in previous studies, and the QIAamp Viral RNA Mini Kit is also used by the WHO for the molecular diagnosis of influenza virus.^1,11,13–15,24–27^

Viral RNA is quantified using molecular methods like quantitative polymerase chain reaction (PCR) and Real-Time qPCR (RT-qPCR). The use of RT-qPCR is considered the gold standard for viral detection and quantification due to the precision, sensitivity and ability to achieve timely results.^29,30^

The purpose of this study is to compare filter types, the effect of air sampling, the use of wash buffers, and RNA extraction methods on virus recovery. The long term goal of this study is to develop optimized methods for performing exposure assessment air sampling targeting influenza virus aerosols. The results of this study will inform future exposure assessment campaigns measuring inhalation exposure to influenza virus.

## Methods

### Fabrication of Polystyrene nanofiber filters

Filters made of PS nanofibers were fabricated via electrospinning, according to the equipment and methodologies described elsewhere.^31^ As detailed in this earlier work, PS filters consisted of nonwoven nanofibers with average diameters of 140 (± 30) nm, a specific surface area (measured by N_2_-BET) of 28 (± 4) m^2^/g, and a water contact angle of 109 ± 9.9, indicative of a hydrophobic surface, as expected.

### Application of Influenza A Virus and Experimental Apparatus

A total of 20 experimental trials were conducted for this experiment. For each trial, two filters of each type of air sampling media (PTFE, PVC, and PS) were spiked with 200 μL of H1N1 influenza virus (strain A/Virginia/ATCC1/2009, ATCC VR-1736, Lot #: 59335953) diluted 1×1000 with HBSS (Ref 14170-112, Lot #: 1227395). The viral solution was allowed to dry on the sampling media inside the biosafety cabinet. A chamber was assembled for each trial to maintain an environment with 50% relative humidity (Figure 1). The chamber functioned as follows: supplied air passed through a HEPA filter (Life Sciences, PN 12144, Lot #: FZ2608) and entered a 1,000 mL flask containing 200-300 mL of molecular grade water, which then entered the chamber. Airflow through the chamber was regulated by a valve attached to an air pump (CRL US, SN: I1001003T). Relative humidity was measured using a direct reading instrument (Extech Instruments Hygro-Thermometer Model SDL500, SN: Z335535) and logged at a rate of one measurement per second. Airflow was adjusted during the experimental trial to maintain a target 50% relative humidity during the entire experiment.

**Figure 1.**
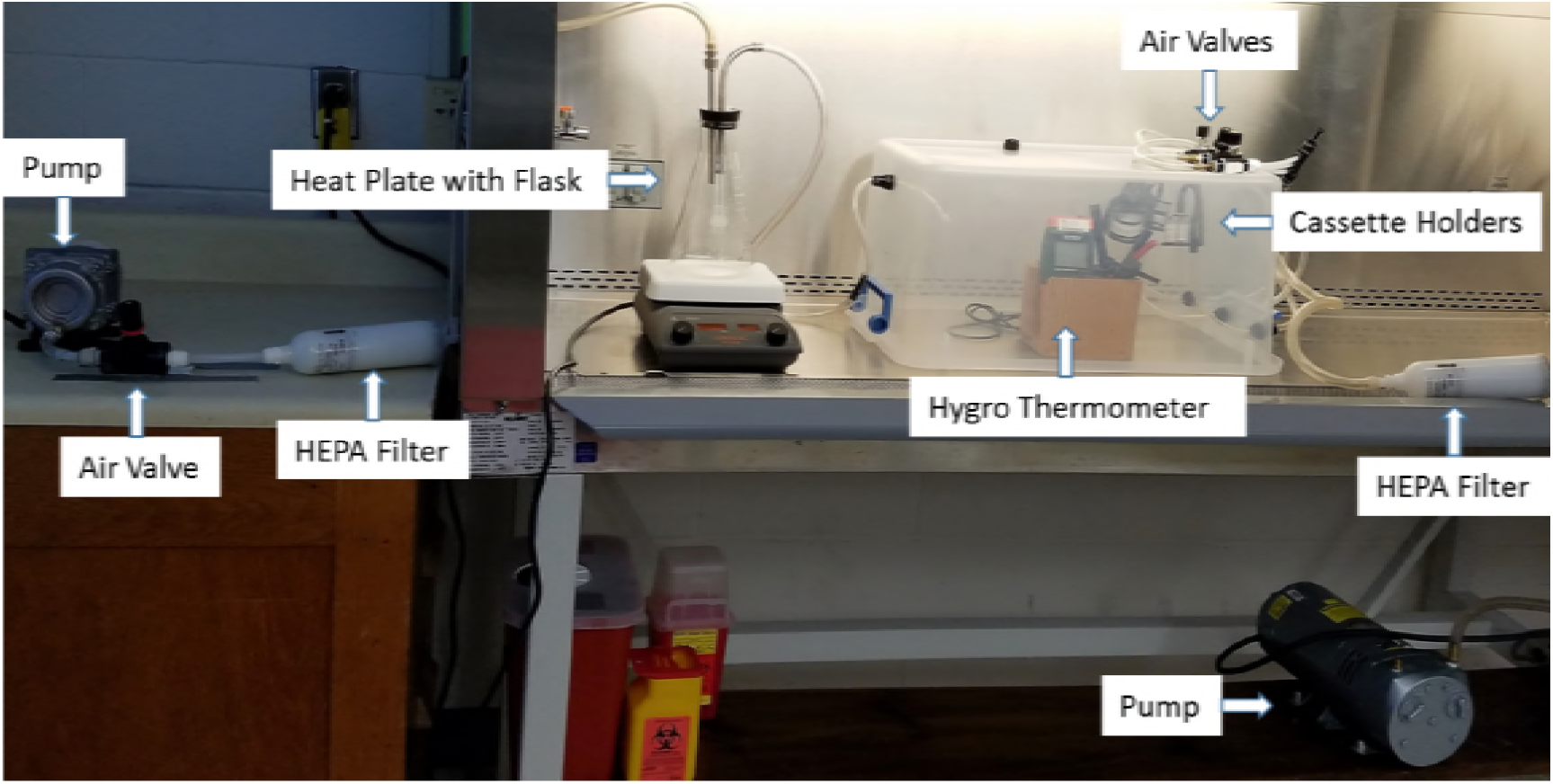
HEPA filtered air was supplied to a flask containing heated water to humidify the air which was then supplied to the sampling chamber where the samples spiked with influenza virus were located.

For every experimental trial, one cassette of each filter material was treated to a simulated air sampling condition of 4 liters per minute (LPM). The tubing for the sampling air, entered in through the right side of the chamber. The sampling train consisted of a cassette holder for each filter type attached with tubing to a GAST Manufacturing vacuum pump (Model No: 0523-101Q-G582DX) with a HEPA filter attached (Life Sciences, PN 12144, Lot #: FZ2608). An airflow control valve (Cole-Parmer Multi-Turn Needle Valve, S/N: 418869-3) was also attached upstream from the pump to control airflow rate. The remaining set of samples were placed at the bottom of the sampling chamber during the experiment to experience similar environmental conditions.

### Sampling Procedure

Once the filters had dried, each individual filter was placed into a 37 mm cassette (SKC, Lot #: 13891-7DD-190) on the surface of a polypropylene backup pad (SKC, Lot #: 13607-7DD-112). The cassettes were sealed with electrical tape, and two cassettes of each filter type were placed into the chamber. One set was treated with sampling air and the remaining set was not treated with sampling air. Calibration of each sampler was performed prior to and after sampling using a primary standard to measure airflow (Bios Defender 520 Model, 717-520M, SKC, USA). Once the Hygro-Thermometer displayed a relative humidity of 50% inside the chamber, the vacuum pump was activated, and the chamber operated for 30 minutes.

After each trial, the data from the Hygro-Thermometer were downloaded, and the average relative humidity was recorded. All filters were placed into 50 mL tubes and washed with 1.5 mL of HBSS or PBS (Cat #: 114-056-101, Lot #: 721749), and then pulse vortexed (Vortex-Genie 2, Scientific Industries, USA) at a speed of 2 for five minutes to remove the virus from the filters. After vortexing, the filters were removed and the tubes were centrifuged at 200 g for two minutes. The sample was aliquoted and stored at −80°C.

### Viral Extraction and Real Time Detection Methodology

Viral RNA was extracted from each sample using a commercial kit [QIAamp Viral Mini RNA Kit (Qiagen, Valencia, CA)] according to the manufacturer’s instructions, along with a Trizol method used in previous literature with some modification.^32^ Superscript® III Platinum One Step RT-qPCR Kit (Life Technologies, Grand Island, NY) was used to synthesize complimentary DNA from 5 uL of viral messenger RNA. The final reaction volume was 25 uL for each sampler. The primers and probe sequences used to complete RT-qPCR were as follows:

- Influenza A forward: 5’- GAC CRA TCC TGT CAC CTC TGA C - 3’
- Influenza A reverse: 5’- AGG GCA TTY TGG ACA AAK CGT CTA - 3’
- Influenza A probe: 6FAM - TGC ACT CCT CGC TCA CTG GGC ACG - BHQ1.

Real Time qPCR used TaqMan reagents (Life Technologies, Grand Island, NY) on a RT-qPCR system (CFX96 Real-Time System, BioRad, USA) for 30 minutes at 50°C, 10 minutes at 95°C, 45 cycles for 15 seconds at 95°C, followed by 35 seconds at 55°C. The maximum cycle number was set to 45 in order to ac QIAGEN for low viral concentrations. All samples were analyzed in triplicates to determine the total amount of virus collected in each sample.

### Calculation of Total Viral Particles

A logarithmic cycle threshold (C_t_) value was determined for each sample by RT-PCR. The C_t_ value is a relative measure of the concentration of the RNA target in the PCR reaction^33^, and is inversely proportional to the original relative expression level of the RNA target.^34^ Using Influenza A plasmid, a standard curve was generated for each RT-qPCR experiment. The antilog (10^x^) was then calculated to get total viral copies in the 5 μL of the isolated RNA PCR reaction. The viral copy data were then volume adjusted to represent the total viral copies per sample. Data were analyzed in R^35^ using multiple linear regression at a single comparison type 1 error rate of 0.05. A random-effects model was also constructed to evaluate evidence for a batch effect, though no systematic batch-to-batch variation was detected.

## Results

Viral recovery for each filter type was significantly different (p-value < 0.0001); with the novel PS filter material recovering the highest Total Viral Copies (TVC), followed by PVC and PTFE, respectively (Tables 1 and 2). Surprisingly, there was not a significant difference in viral RNA among filters treated with air compared to filters not treated with air (Kit, HBSS: Air vs. No-Air p-value = 0.615; Trizol, HBSS: Air vs. No-Air p-value = 0.947; Kit, PBS: Air vs. No-Air p-value = 0.224; Trizol, PBS: Air vs. No-Air p-value = 0.112). TVCs were significantly different among samples using HBSS and PBS as filter wash buffers, however the difference was dependent upon the extraction method used (Kit, Air or No-Air: HBSS vs. PBS p-value = 0.0001, 0.0001; Trizol, Air or No-Air: HBSS vs. PBS p-value = 0.0322, 0.499). HBSS showed greater viral recovery than PBS when using HBSS, but slightly lower recovery when using Trizol. TVCs were significantly different when using the Kit and Trizol extraction methods for all comparisons (HBSS, Air or No-Air: Kit vs. Trizol p-value = 0.0021, 0.0013; PBS, Air or No-Air: Kit vs. Trizol p-value = 0.0001, 0.0002). TVCs were higher with the Kit when using HBSS, and higher with the Trizol method when using PBS (Figure 2).

**Table 1.** Geometric Mean (GSD) Total Influenza Viral Copies: With Air

| Extraction | Wash | Filter Type |  |  |
| --- | --- | --- | --- | --- |
|  |  | PTFE<sup>†</sup> | PVC<sup>†</sup> | PS |
| Kit | PBS | 158 (4) | 626 (7) | 14,240<sup>†</sup> (7) |
|  | HBSS | 1,755<sup>‡</sup> (3) | 24,155<sup>‡</sup> (3) | 179,442<sup>‡†</sup> (1) |
| Trizol | PBS | 1,898<sup>‡</sup> (3) | 13,954<sup>‡</sup> (1) | 57,176<sup>‡†</sup> (2) |
|  | HBSS | 564 (4) | 8,325 (4) | 41,164<sup>†</sup> (5) |

<sup>†</sup> Comparison across filter types ( $p$ -value = 0.0001)
<sup>‡</sup> Comparison across wash buffers, within extraction method ( $p$ -value = 0.0001)

**Table 2.** Geometric Mean (GSD) Total Influenza Viral Copies: Without Air

| Extraction | Wash | Filter Type |  |  |
| --- | --- | --- | --- | --- |
|  |  | PTFE<sup>†</sup> | PVC<sup>†</sup> | PS |
| Kit | PBS | 117 (5) | 755 (8) | 5,081<sup>†</sup> (3) |
|  | HBSS | 2,599<sup>‡</sup> (2) | 29,220<sup>‡</sup> (2) | 162,897<sup>†‡</sup> (2) |
| Trizol | PBS | 726<sup>‡</sup> (3) | 9,939<sup>‡</sup> (2) | 51,020<sup>†‡</sup> (1) |
|  | HBSS | 537 (4) | 9,057 (5) | 39,942<sup>†</sup> (4) |

<sup>†</sup> Comparison across filter types (p-value = 0.0001)
<sup>‡</sup> Comparison across wash buffers, within extraction method (p-value = 0.0001)

**Figure 2.**
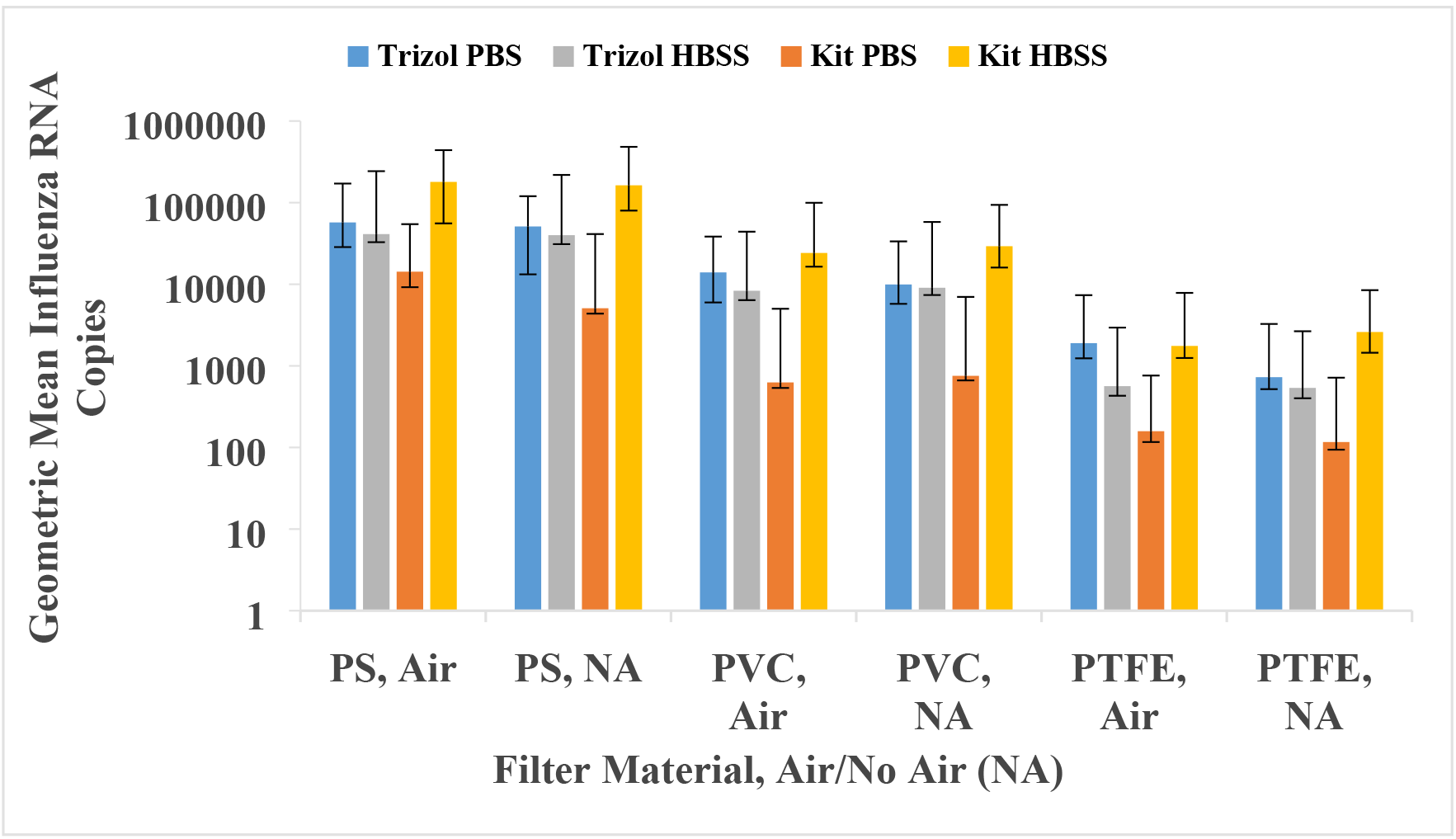
Geometric mean influenza RNA copies and standard deviation for each condition: Air or No air, HBSS or PBS wash buffer, and Trizol or Kit extraction method. Y-axis is a logarithmic scale. Error bars represent the geometric standard deviation.

## Discussion

Viral recovery was significantly different for each filter type (p-value < .0001); with the novel electrospun PS filter material resulting in the recovery of the greatest amount of viral RNA, followed by PVC and PTFE. PTFE is often considered the optimal filter material in virus collection^10^; however, the results from this experiment suggest that PS outperforms PTFE for viral recovery. To the best of our knowledge, no previous studies have evaluated the RNA viral recovery from PS used as a filter material in field of exposure science.

The underlying cause for the superior performance of PS relative to PTFE and PVC merits further investigation. There are two notable differences in the PS relative to the other materials used in this experiment. First, the PS filter material was fabricated differently than the other filter materials. Electrospinning produces a nanostructured mat consisting of nonwoven fibers, whereas the other polymers (PTFE and PVC) are more traditional polymer films. The nanofiber mat of PS is sufficiently dense and may capture a greater number of viruses and release them efficiently compared to PTFE and PVC polymer films. Additional work is needed to directly compare PS nanofiber mats to PTFE or PVC also fabricated by electrospinning, thus producing similarly nanostructured layers.

Alternatively, rather than just physical differences, differences in the chemical composition of the polymers may result in the increased performance. Specifically, the base monomers of PTFE [(C_2_F_4_)_*n*_] and PVC [(C_2_H_3_Cl)_*n*_] contain halogen atoms (fluorine and chlorine, respectively), whereas PS [(C_8_H_8_)_*n*_] is an aromatic hydrocarbon-based polymer. Carbon-halogen bonds are more polar, and thus potentially more reactive, than simple hydrocarbon bonds.^36^ Thus, chemical environments produced by surface fluorine and chlorine may result in conditions that affect the viability of RNA captured on more traditional, halogen-containing polymer materials (*i.e.*, PTFE and PVC).^37^

In this experiment, we failed to detect evidence that the treatment of filters with simulated sampling air affected the recovery of RNA from the filter materials (Kit, HBSS: Air vs. No-Air p-value = 0.615; Trizol, HBSS: Air vs. No-Air p-value = 0.947; Kit, PBS: Air vs. No-Air p-value = 0.224; Trizol, PBS: Air vs. No-Air p-value = 0.1122). Due to the negative affect that filter materials have had on viral recovery^10,19^ it was hypothesized that air being pulled through filter materials would affect the viral recovery. The results of this experiment dispute this observation given the environmental and sampling conditions used in this experiment are similar previous studies using air sampling.^9,25^

Using HBSS and PBS as a filter wash buffer resulted in viral recoveries that were significantly different, depending on the RNA extraction method used (Kit, Air or No-Air: HBSS vs. PBS p-value = 0.0001, 0.0001; Trizol, Air or No-Air: HBSS vs. PBS p-value = 0.0322, 0. 499). Using HBSS as a wash buffer resulted in greater viral recovery then PBS when using the QIAamp Viral RNA Mini Kit, but slightly lower recovery when using HBSS and Trizol. Our results align with previous studies which indicate that HBSS and PBS can be used in the removal of virus from sampling materials^13,14,17^, however no other studies have looked at the effect different wash buffer has on viral recovery.

Viral RNA copies were significantly different when using the Kit and Trizol RNA extraction methods for all comparisons (HBSS, Air or No-Air: Kit vs. Trizol p-value = 0.0021, 0.0013; PBS, Air or No-Air: Kit vs. Trizol p-value = 0.0001, 0.0002). Viral RNA copies were higher with the QIAamp Viral RNA Mini Kit when using HBSS, and higher with the Trizol method when using PBS. Previous studies have found that Trizol extraction methods typically produce higher copies of viral RNA when compared to other extraction methods^24,38,39^, however previous studies have not evaluated whether wash buffer type affects viral RNA copies as was observed in this study.

### Limitations

In this experiment, viral RNA copies for the extraction processes varied considerably at times among experimental conditions. For the Trizol method, a visible RNA “pellet” was to be observed after extraction, distinguishable from impurities in the sample. However, several samples did not produce a visible pellet, making it more difficult to distinguish between the viral RNA and the remaining impurities in the sample. Therefore, the Trizol method may have resulted in additional loss of viral RNA. Viral recovery efficiency from each filter type was not measured for each sample; therefore, it is unknown whether the differences in virus recovered from the filter types in each sample had an effect on viral RNA copy variability.

After washing the filters with either HBSS or PBS, aliquots for each extraction process were separated into individual conical tubes. Although samples were pulse vortexed in an effort to provide an even distribution of virus in the solution, the virus may not have been distributed evenly during the preparation of aliquots. This error would affect the amount of viral RNA extracted in each aliquot, biasing the TVC of one extraction process, wash buffer, or filter type over the other. However, we expect this error to be non-differential in this experiment.

Viral losses may have occurred throughout this experiment. Pipetting errors could have occurred while applying viral solution to filter materials, transferring virus to multiple conical tubes, and during extraction processes. Viral losses may have also occurred during the multiple freeze thaw cycles of each sample, and during RT-qPCR cycles. However, we expect viral losses to be non-differential in this experiment.

Viral activity was not measured because RT-qPCR is unable to provide information on virus activity. Determining whether virus particles collected are active (i.e., viable and contagious) is important in the prevention and control of influenza virus infection. Therefore, infectivity assays or other methods to assess the viral activity are needed and should be included in future studies.

For this experiment, a target of 50% RH was established for all trials. Although the optimal comfort level for humans is 40-60% RH^14^, and previous studies have found that higher RHs (i.e., 50-56%) have no effect on viral infectivity.^13^ Relative humidity was measured using a single real time measurement instrument in the chamber over the course of the experiment. We assumed the experimental chamber was well mixed however, the RH may not have remained at 50% for all sampling locations during the entire 30-minute sampling period.

Filters were spiked with a known concentration of influenza virus solution in a controlled laboratory setting. As mentioned previously, environmental factors like temperature and RH can affect viral recovery^10^, and a major route of influenza virus transmission is through aerosolized virus particles.^5,6^ Therefore, the results observed in this study may not be generalizable for all environmental conditions.

## Conclusions

Viral recovery for each filter type was significantly different (p-value < 0.0001), with the novel electrospun PS filter material resulted in the recovery of the most viral RNA, followed by PVC and PTFE. Treating filters with simulated sampling air did not significantly affect the recovery of RNA from the filter materials (Kit, HBSS: Air vs. No-Air p-value = 0.615; Trizol, HBSS: Air vs. No-Air p-value = 0.947; Kit, PBS: Air vs. No-Air p-value = 0.224; Trizol, PBS: Air vs. No-Air p-value = 0.1122). Future studies should attempt to aerosolize virus at concentrations similar to environmental concentrations, to determine if using the novel PS filter material will result in detection and quantification of influenza virus. In addition, future studies should compare the viral recovery of PS filter material with other commonly used samplers that do not use filter material as a sampling media (i.e., SKC Biosampler and Anderson N6 impactor).

Using HBSS and PBS as a filter wash buffer resulted in viral recoveries that were significantly different, depending on the RNA extraction method used (Kit, Air or No-Air: HBSS vs. PBS p-value = 0.0001, 0.0001; Trizol, Air or No-Air: HBSS vs. PBS p-value = 0.0322, 0.499). Using HBSS as a filter wash buffer resulted in greater viral recovery than PBS when using HBSS, but slightly lower recovery when using Trizol. Previous studies have not compared viral recovery and quantification between HBSS and PBS. Viral RNA copies were significantly different when using the Kit and Trizol RNA extraction methods for all comparisons (HBSS, Air or No-Air: Kit vs. Trizol p-value = 0.0021, 0.0013; PBS, Air or No-Air: Kit vs. Trizol p-value = 0.0001, 0.0002). Viral RNA copies were higher with the Kit when using HBSS, and higher with the Trizol method when using PBS. Future studies examining exposure to influenza virus aerosols should vary the filter wash buffer base on the approach used for RNA extraction. Also future studies should consider using PS as a filter media for aerosol sampling targeting influenza virus.

## Acknowledgments

Financial aid and equipment support was provided by the Heartland Center for Occupational Health and Safety, a NIOSH/CDC Education and Research Center (T420H008491).

